# Layer-5 Pyramidal Cell tLTD Requires Astrocytic Ca^2+^ and CB1 Receptor Signaling

**DOI:** 10.64898/2026.04.21.719669

**Authors:** Airi Watanabe (渡辺愛利), Connie Guo, Shawniya Alageswaran, P. Jesper Sjöström

## Abstract

Timing-dependent long-term depression (tLTD) is a form of spike timing-dependent plasticity (STDP) that has been attributed to presynaptic NMDA and CB1 receptors at synapses between layer 5 (L5) pyramidal cells (PCs) in visual cortex. Here, we asked whether astrocytes, known for gliotransmission and tripartite synapse formation, are also required. Using quadruple whole-cell recordings in acute slices from C57BL/6 mice, we found that L5 PC → PC tLTD was abolished by the glial metabolic inhibitor sodium fluoroacetate. Disrupting astrocyte Ca^2+^ signaling through loss-of-function approaches such as AAV-mediated expression of CalEx and BAPTA loading of astrocyte networks consistently prevented tLTD. Optogenetic activation of astrocyte Gq signaling during tLTD induction abolished tLTD and led to potentiation. Since tLTD relies on endocannabinoid (eCB) signaling and astrocytes express CB1 receptors, we conditionally deleted CB1 receptors from astrocytes and found that this manipulation abolished tLTD. Taken together, our results show that L5 PC → PC tLTD requires astrocyte Ca^2+^ signaling and CB1 receptor activation. These findings suggest that astrocyte-dependent control of STDP may represent a general principle across circuit and synapse types.

**Significance Statement:** Spike timing-dependent plasticity (STDP) enables neural circuits to adapt based on the precise timing of activity, with timing-dependent long-term depression (tLTD) promoting competition and stabilization. While tLTD is often considered a two-factor process involving pre- and postsynaptic neurons, we show that astrocytes form a key third element. Using targeted loss-of-function approaches, we demonstrate that tLTD at layer-5 pyramidal cell synapses in visual cortex requires astrocyte Ca^2+^ signaling and CB1 receptors. Unexpectedly, optogenetic stimulation of astrocyte Gq signaling during induction blocked depression and instead triggered potentiation. These findings call for a revision of existing models of STDP and suggest that astrocyte-dependent plasticity may reflect a general regulatory principle across cortical circuits.

## Introduction

Synaptic plasticity is considered a cornerstone of memory formation (Bliss and Collingridge, 1993; Malenka and Bear, 2004; Nabavi et al., 2014) and a key mechanism in the developmental refinement of neural circuits (Katz and Shatz, 1996; Cline, 1998; Song and Abbott, 2001). This foundational idea is often traced back to Donald Hebb (1949) and is captured by the slogan “cells that fire together wire together” (Shatz, 1992), which highlights the role of correlated activity of connected neurons in strengthening synapses.

In more recent years, researchers have emphasized the role of temporal ordering of neuronal activity in shaping synaptic changes, a process known as spike timing-dependent plasticity, or STDP (Feldman, 2012; Markram et al., 2012). Early studies revealed that when a presynaptic neuron fires within a ten-millisecond before a postsynaptic partner, the connection is strengthened through timing-dependent long-term potentiation (tLTP), but when the order is reversed, the connection weakens through timing-dependent long-term depression (tLTD) (Markram et al., 1997; Bi and Poo, 1998; Zhang et al., 1998; Feldman, 2000; Sjöström et al., 2001). This bidirectional form of plasticity thus extends Hebb’s original idea by incorporating tLTD, which enables key computational features such as synaptic competition (Song et al., 2000) and network stabilization (Song and Abbott, 2001).

Both classical Hebbian plasticity and STDP are examples of two-factor learning rules, meaning they depend on coordinated activity in presynaptic and postsynaptic neurons (McFarlan et al., 2023). However, research over the past couple of decades have highlighted that other cells — including astrocytes in a tripartite synaptic structure (Perea et al., 2009) — can also contribute to long-term plasticity at specific synapse types (Henneberger et al., 2010; Valtcheva and Venance, 2016; Adamsky et al., 2018).

We previously demonstrated that tLTD at synapses between visual cortex L5 PCs requires simultaneous activation of endocannabinoid CB1 receptors and presynaptic NMDA receptors (Sjöström et al., 2003). Based on this, we proposed a model in which both receptor types are located at release sites in L5 PC axons (Sjöström et al., 2003; Duguid and Sjöström, 2006). However, more recent work at neocortical L4 → L2/3 synapses (Min and Nevian, 2012) as well as at hippocampal CA3 → CA1 synapses (Andrade-Talavera et al., 2016; Falcón-Moya et al., 2020) demonstrated that tLTD relies on CB1 receptor signaling in astrocytes. This raised the possibility that at L5 PC → L5 PC connections, the CB1 receptors relevant for tLTD are not those located in L5 PC axons (Katona et al., 2006) but those in nearby astrocytes (Busquets-Garcia et al., 2018).

We therefore revisited our original model of L5 PC → L5 PC tLTD (Sjöström et al., 2003; Duguid and Sjöström, 2006), testing whether astrocyte signaling is also required. Using a range of targeted interventions, we found that astrocytes are in fact necessary for the induction of L5 PC → PC tLTD. Specifically, we found that astrocyte signaling via Ca^2+^, Gq protein, and cannabinoid CB1 receptors critically determines L5 PC → L5 PC tLTD. Our findings suggests that astrocyte-mediated control of STDP may be a general principle that holds across circuit and synapse types.

## Methods

### Animals and Ethics Statement

The animal study was reviewed and approved by the Montreal General Hospital Facility Animal Care Committee and adhered to the guidelines of the Canadian Council on Animal Care. Mice were kept on a 12h light and 12h dark cycle, provided with enrichment such as bio-huts and chew blocks, and given food and water *ad libitum*.

C57BL/6J mice (referred to as WT here) were obtained from The Jackson Laboratory (JAX strain 000664). Hemizygous Aldh1ll-Cre mice (Tien et al., 2012) were obtained from The Jackson Laboratory (JAX 023748) and maintained by crossing to WT animals. Homozygous Emx1-Cre mice (Gorski et al., 2002) were also obtained from The Jackson Laboratory (005628). Cryorecovered heterozygous CB1^f^ mice (JAX 036107) were used to conditionally delete the cannabinoid receptor 1 *Cnr1* gene (Marsicano et al., 2003). *Cnr1*^fl/+^ mice that were cryorecovered were crossed to obtain *Cnr1*^fl/fl^ mice.

To determine mouse genotypes, tail biopsy and paw tattooing were performed prior to postnatal day 6 (P6). Genomic DNA was extracted using standard methods, and genotyping was carried out using Jackson Laboratory-recommended primers (Aldh1l1-Cre: 14314, 18707, 18708; CB1f: 57165, 57166). PCR reactions were performed using HotStarTaq DNA Polymerase kit (QIAGEN, 203203) and dNTPs from Invitrogen/Thermo Fisher (18427–013).

### Acute Slice Preparation

Male and female mice aged P11 – P28 were anesthetized with isoflurane and swiftly decapitated once the hind-limb withdrawal reflex was lost. The brain was rapidly extracted and submerged in ice-cold (<4°C) artificial cerebrospinal fluid (ACSF), containing in mM: 125 NaCl, 2.5 KCl, 1.25 NaH_2_PO_4_, 26 NaHCO_3_, 25 glucose, 1 MgCl_2_ and 2 CaCl_2_. ACSF solution was constantly bubbled with 95% O_2_/5% CO_2_ and osmolality was adjusted to 338 mOsm, measured using Model 3300, Model 3320 or Osmo1 osmometers (Advanced Instruments Inc., Norwood, MA, USA). Oblique coronal 300 µm-thick slices were cut using a Campden Instruments 5000 mz-2 vibratome (Lafayette Instrument, Lafayette, IN, USA). Slices were incubated at ∼33°C in ACSF for 15–30 min, then allowed to recover at room temperature (∼23°C) for at least one hour after slicing before recording.

For mice aged ∼P16 to ∼P21, slices were sectioned in low-Ca^2+^ ACSF (2 mM MgCl_2_, 1 mM CaCl_2_). For mice ∼P21 and older, slicing was performed in sucrose-based solution containing (in mM): 200 sucrose, 2.5 KCl, 1.0 NaH_2_PO_4_, 2.5 CaCl_2_, 1.3 MgCl_2_, 47 glucose, and 26.2 NaHCO_3_. These slices were also recovered in low-Ca^2+^ ACSF prior to recording in standard ACSF as described above.

### Electrophysiology

We carried out experiments with ACSF heated to 33°C (resistive inline heater, Scientifica Ltd, Uckfield, UK; or TC-324C + SH-27B, Warner Instruments, Holliston, MA, USA). Temperature was continuously recorded and verified offline, and recordings were truncated or excluded if outside the 32°C to 34°C range.

Patch pipettes (typically 4 – 7 MΩ) were pulled from fire-polished glass capillaries (BF150-86-10, Sutter Instruments, Novato, CA, USA) using a P-1000 puller (Sutter Instruments, Novato, CA, USA) and filled with internal solution (in mM: KCl, 5; K-Gluconate, 115; HEPES, 10; Mg-ATP, 4; Na-GTP, 0.3; Na-Phosphocreatine, 10; Biocytin, 0.1% w/v; adjust with KOH to pH 7.2; adjusted with sucrose to 310 mOsm).

Depending on the experiment, the internal solution also included one of the following: Alexa Fluor 594 (15–40 µM; A10428, Life Technologies), Alexa Fluor 488 (20–100 µM; A10436, Life Technologies), BAPTA (20 mM; B1204, Fisher Scientific), or Fluo-5F (200 µM; F14221, Fisher Scientific).

Whole-cell recordings were obtained using either BVC-700A (Dagan Corporation, Minneapolis, MN, USA) or Model 2400 (A-M Systems, Carlsborg, WA, USA) amplifiers, with signals filtered at 5 kHz and sampled at 40 kHz using PCIe-6323 boards (NI, Austin, TX, USA) controlled by MultiPatch custom software (https://github.com/pj-sjostrom/MultiPatch.git) running in Igor Pro 8 or 9 (WaveMetrics Inc., Lake Oswego, OR, USA).

L5 PCs were targeted for patching based on their large and characteristically pyramid-shaped somata. Cell identity was further assessed via a series of somatic current injections (500 ms; –0.3 to +0.7 nA in 0.2 nA steps), which evoked spiking patterns typical for L5 PCs. Morphological confirmation was obtained post hoc using two-photon (2p) imaging (see below). Astrocytes were identified by preincubating slices in ACSF containing sulforhodamine 101 (1 µM; S7635, Millipore Sigma, ON, Canada) for 5 minutes at room temperature. SR101-labeled astrocytes were then visualized by 2p microscopy and targeted for whole-cell recording and/or imaging. Astrocytes were loaded with Fluo-5F for at least 15 min before Ca^2+^ imaging.

To find monosynaptically connected pairs among L5 PCs, for which connectivity is only a sparse 10 – 15% (Song et al., 2005; Chou et al., 2024), we employed quadruple whole-cell recordings to test up to twelve candidate connections simultaneously. Four neurons were identified and approached with four pipettes. Gigaohm seals were formed sequentially, followed by successive break-in. To screen for synaptic connections, five spikes were evoked at 30 Hz using 1.3 nA current injections in each cell, repeated up to 20 times at 20-second inter-sweep intervals. In a subset of experiments, synaptic responses were evoked using either extracellular stimulation or a targeted optogenetic method described below (optomapping). For extracellular stimulation, the pipette was filled with filtered ACSF and placed approximately 100 µm from the patched cell to recruit local inputs.

### Plasticity Protocols

Prior to drug application or plasticity induction, a baseline period of 7–20 minutes was recorded. During this period, bursts of 5 spikes were evoked using trains of 5-ms-long ∼1.3-nA current pulses delivered at 30 Hz every 20 seconds. We induced tLTD using a post-before-pre spike pairing protocol consisting of five postsynaptic spikes at 20 Hz, timed either 10 ms or 25 ms before presynaptic stimulation, repeated 15 times with 10-second intervals. This protocol is known to reliably elicit tLTD at L5 PC → PC synapses (Sjöström et al., 2003). Following the induction, the baseline 30-Hz spike pattern was resumed for up to 75 minutes. All recordings were performed in current clamp. A subset of plasticity experiments (Fig. 3) were carried out with 2p optogenetic activation (Chou et al., 2024), as previously described (Chou et al., 2025), see below for additional details.

Recordings with unstable baselines were excluded based on a t-test of Pearson’s r for response amplitudes (Lalanne et al., 2016). Input and series resistance were assessed using a 250-ms-long hyperpolarizing test pulse of -25 pA. Recordings were excluded or truncated if input resistance changed by more than 30%, or if resting membrane potential shifted by more than 8 mV. Recordings shorter than 20 minutes post induction or drug wash-in were excluded from analysis.

The locus of plasticity, i.e., whether plasticity was expressed pre- or postsynaptically, was assessed using paired-pulse ratio (PPR) and coefficient of variation (CV) analyses (Brock et al., 2020). PPR was calculated as the ratio of the second EPSP to the first, or EPSP_2_ / EPSP_1_. The change in PPR, or ΔPPR, was computed by subtracting PPR measured after drug wash-in or plasticity induction from the baseline PPR, or PPR_after_ - PPR_before_. If the presynaptic cell spiked twice for individual 5-ms-long current pulses, the experiment was excluded from PPR analysis. For PPR and CV analyses, experiments with EPSPs weaker than 0.3 mV were excluded, to ensure sufficient signal to noise.

### Neonatal Viral Injection

All adeno-associated viruses (AAVs) were aliquoted on arrival and stored at –80 °C until use. The following constructs were used for neonatal intracerebral injections: AAV2/5-GfaABC1D-mCherry-hPMCA2w/b (Addgene 111568-AAV5), AAV9-CAG-DIO-ChroME-ST-P2A-H2B-mRuby3 (Addgene 108912-AAV9), AAV1-GfaABC1D-OptoGq-eYFP (Adamsky et al., 2018), AAV5-GfaABC1D-Cre-4×6T (Addgene 196410-AAV5), and AAV5-pCAG-FLEX-EGFP-WPRE (Addgene 51502-AAV5).

The following viral titers were used (in genome copies per milliliter, GC/ml): 1.20×10^13^ for hPMCA2w/b, 2.7×10^12^ for ChroME, 1.60×10^12^ to 1.60×10^13^ for OptoGq, 1.60×10^13^ for GfaABC1D-Cre, and 1.50×10^13^ for FLEX-EGFP.

Intracerebral injections were performed in P0–P2 mice, targeting visual cortex (Kim et al., 2014; Chou et al., 2024). Mice were cryoanesthetized by placing them on aluminum foil over ice for approximately 10 minutes, then secured in a neonatal stereotaxic apparatus. Injections were delivered using a 33-gauge needle (point style 4, 30° bevel, 25.4 mm length) mounted on a 10 µl gas-tight syringe (Hamilton Instruments, Reno, NV, USA), controlled by a syringe pump (Model 70-4507, Harvard Apparatus, Holliston, MA, USA).

The injection site was located at 0.00 mm anterior-posterior and 1.1 mm medial-lateral relative to lambda. Injections were delivered at three depths along a single needle track (0.2, 0.15, and 0.1 mm below the pial surface) during stepwise retraction of the needle, at a constant rate of 0.25 µl/min. At each depth, volumes of 0.35 µl were infused for hPMCA2w/b, GfaABC1D-Cre, and FLEX-EGFP. For ChroME, 0.3–0.4 µl was injected; for OptoGq, 0.2–0.4 µl was used. Following injection, pups were placed under a heat lamp and monitored until spontaneous movement resumed.

### Two-Photon Imaging

We used custom-built imaging workstations for 2p imaging, as previously described (Abrahamsson et al., 2017), using either a Chameleon Ultra II (Coherent, PA, USA) or a Mai Tai HP (Spectra-Physics, CA, USA) tunable Ti:Sa laser. Lasers were tuned to wavelengths optimal for visualization of the dye at hand, e.g., 750 nm for mCherry and Alexa 488, 820 nm for Alexa-594, 1040 nm for mRuby, 920 nm for eYFP, etc.

Laser power was monitored using a power meter and controlled via a half-wave plate and polarizing beam-splitting cube (Thorlabs GL10-B and AHWP05M-980). Gating was achieved with a mechanical shutter (Thorlabs SH05/SC10), and scanning was performed using 6215H galvanometric mirrors (3 mm; Cambridge Technology, Bedford, MA, USA).

Fluorescence was detected using bialkali photomultipliers (2PIMS-2000-20-20, Scientifica Ltd., Uckfield, UK) through an Olympus LUMPlanFL N 40×/0.80 objective, or via substage-mounted R3896 photomultipliers (Hamamatsu, Bridgewater, NJ, USA) collected through an Olympus Achromatic/Aplanatic condenser (NA 1.40). Emission light was split using a Semrock FF665 dichroic, and laser light was blocked with a Semrock FF01-680 filter. Red and green fluorescence channels were separated with a Chroma t565lpxr dichroic in combination with Chroma ET630/75M and ET525/50M emission filters.

Laser-scanning Dodt contrast was generated using custom optics by collecting transmitted laser light through the acute slice with a spatial filter and diffuser placed in a 1× telescope, focused onto an amplified photodiode (Thorlabs PDA100A-EC). Signals from photomultipliers and the photodiode were digitized using PCIe-6374 boards (National Instruments, Austin, TX, USA) and acquired with ScanImage (2019–2024) running in MATLAB (MathWorks, Natick, MA, USA).

Astrocyte Ca^2+^ activity was imaged using 2p excitation at 780 – 930 nm to visualize changes in Fluo-5F fluorescence. Each movie was taken at a single focal plane for 150 seconds using 512 × 512-pixel frames acquired at 2.11 Hz. Movies were taken before and after drug wash-in or tLTD induction. Ca^2+^ signals were measured as dG/R, a change in green Fluo-5F fluorescence normalized to red SR101 fluorescence. For the detection of Ca^2+^ events, at least 10 regions of interests (ROIs) of were manually selected and baseline was set at the 10 – 20 frames with the weakest Ca^2+^ signal. We low-pass filtered at 0.2 Hz to eliminate high-frequency noise. Ca^2+^ events were detected by a simple thresholding algorithm set to detect signal at 0.5 – 1 sigma above background noise (Watanabe et al., 2023). Mean Z-scores of Pearson’s r for the correlation of all ROIs were used to measure the correlation of Ca^2+^ activity within cells (Watanabe et al., 2023). Astrocytes that depolarized to >-40 mV during imaging, cells with low signal to noise ratio whereby at least 10 ROIs could not be selected, and movies with significant drift that could not be corrected for (e.g., Z-plane drift) were excluded from analysis. Imaging data were analyzed using LineScan Analysis (https://github.com/pj-sjostrom/LineScanAnalysis) running in Igor Pro.

### Optogenetics

We used 2p optogenetic stimulation, or optomapping (Chou et al., 2024), to recruit presynaptic cells onto a patched postsynaptic PC in acute slices from Emx1^Cre/Cre^ mice injected with AAV-ChroME. A Mai Tai HP Ti:Sa laser tuned to 1040 nm was used to evoke action potentials in ChroME-expressing neurons via spiral scanning with jScan software (https://github.com/pj-sjostrom/jScan) running in Igor Pro 9. Each opsin-expressing candidate cell was stimulated with two spiral scans 500 ms apart, while monitoring the postsynaptic response.

To exclude directly activated postsynaptic PCs, responses with onset latency <1 ms were discarded (Chou et al., 2024). Upon identifying a synaptic connection, only the connected presynaptic cell was repeatedly stimulated to record a ∼10 min baseline, induce tLTD, and record a post-induction period, following electrophysiology parameters described above. Responses <0.3 mV EPSP amplitude were excluded. The power of the Ti:Sa laser was set to ∼55 mW as measured at the objective back aperture.

To activate the light-sensitive Gq-coupled receptor Opto-α1AR in C57BL/6J mice injected with OptoGq virus, we used 445-nm laser light (Industrial 600 mW Blue Laser Module, eBay.ca, Item ID: 181041391795). Light was delivered via the 40× objective in 45 ms pulses at 20 Hz during tLTD induction. The 445-nm beam was combined with the 2p laser using a Semrock FF665 dichroic and controlled via the MultiPatch software to allow synchronized stimulation with electrophysiology recordings. The power of the 445-nm laser was set to ∼13 mW as measured at the objective back aperture.

### Pharmacology

Acute slices were preincubated in ACSF containing 5 mM sodium fluoroacetate (NaFAC; MP Biomedicals, CA, USA) for 30 min prior to recordings; NaFAC was not present during experiments. Arachidonyl-2-chloroethylamide (ACEA; Cayman Chemical, MI, USA) was used at 125 nM in ACSF and was washed in following baseline acquisition.

### Biocytin Staining and Confocal Imaging

To preserve patched neurons for biocytin histology, patch pipettes were slowly retracted after recordings to allow resealing of the membrane. Acute slices were fixed overnight at 4 °C in 4% paraformaldehyde and then transferred to 0.01 M phosphate-buffered saline (PBS) for storage (up to 2 weeks) prior to staining.

For biocytin-only staining, slices were washed 4× for 10 min each in 0.01 M Tris-buffered saline (TBS) with 0.3% Triton X-100, blocked for 1 h in the same buffer containing 10% normal donkey serum (NDS; 017-000-121, Jackson ImmunoResearch), then incubated overnight at 4 °C with 1:200 Alexa Fluor 647–conjugated streptavidin (S32357, ThermoFisher Scientific) in TBS with 0.3% Triton X-100 and 1% NDS. Slices were then washed 4× for 10 min in TBS.

For combined biocytin and immunofluorescence staining, slices were washed 4× for 5 min in 0.01 M PBS with 1% Triton X-100, blocked for 1.5 h in PBS with 0.3% Triton X-100 and 10% NDS, and incubated overnight at 4 °C with 1:500 Alexa Fluor 647–streptavidin and 1:1000 monoclonal rat anti-mCherry antibody (M11217, Invitrogen) in PBS containing 1% NDS and 0.3% Triton X-100. The next day, slices were washed 3× in the same buffer and incubated for 2 h with 1:1000 Alexa Fluor 488–conjugated donkey anti-rat secondary antibody (712-585-150, Jackson ImmunoResearch), followed by three additional washes. Slices were mounted on SuperFrost Plus slides (1255015, ThermoFisher) using ProLong Glass Antifade Mountant (P36984, ThermoFisher), and the edges of the coverslip were sealed with clear nail polish after curing.

Image stacks were acquired using Zeiss LSM 780 or 880 confocal microscopes at 10× or 20× magnification with ZEN2010 software (ZEISS International). We used the following laser lines: 633 nm for Alexa Fluor 647, 561 nm for mRuby, and 488 nm for Alexa Fluor 488 and for EGFP.

### Statistics

Unless otherwise noted, results are reported as mean ± standard error of the mean (SEM). Boxplots show medians and quartiles, with whiskers denoting extrema. Significance levels are indicated as * p < 0.05, ** p < 0.01, and *** p < 0.001.

For multiple comparisons, pairwise testing was performed only if ANOVA indicated significance at the p < 0.05 level. Standard ANOVA was used for homoscedastic data and Welch’s ANOVA for heteroscedastic data, based on Bartlett’s test (p < 0.05). Pairwise comparisons employed two-tailed Student’s t-tests for equal means. If an F-test for equality of variances indicated p < 0.05, the unequal variances t-test was used instead. Bonferroni’s correction was always used to adjust for multiple comparisons. For data that was not normally distributed (e.g., Supp. Fig. 2B), we relied on the Kruskal-Wallis test instead of ANOVA, and Mann-Whitney U test instead of Student’s t-test.

For CV analysis, φ was defined as the angle relative to the diagonal demarcation line and Wilcoxon signed rank test was used to compare φ relative to the diagonal (Brock et al., 2020). Changes in 1⁄𝐶𝑉^2^_*norm*_ were compared using a one-sample Student’s t test versus 1.

Statistical tests were carried out in Igor Pro 9 (WaveMetrics Inc., OR, USA). Linear mixed-effects models were implemented in RStudio 2024.12.0+467 and 2025.09.2+418 (Posit Software, Boston, MA, USA), with Tukey’s method for post hoc adjustments. For the mixed-effects models, age and genotype were treated as fixed effects, and cell as random effect. To account for log-normal synaptic strength distribution (Chou et al., 2024), EPSP amplitudes were log-transformed prior to parametric statistical comparisons.

## Results

### Astrocytes are involved in L5 PC → PC tLTD

To test whether astrocytes contribute to L5 PC → PC tLTD, we preincubated acute visual cortex slices with the glial metabolic inhibitor NaFAC (Swanson and Graham, 1994). Because L5 PC → PC connectivity rate is low (∼10-15%, Song et al., 2005; Chou et al., 2024), we used quadruple whole-cell recordings (Fig. 1A) to find monosynaptic connections (Lalanne et al., 2016). In control slices, L5 PC → PC connections reliably underwent tLTD. However, in the presence of NaFAC, tLTD was abolished (Fig. 1B–E).

**Figure 1.**
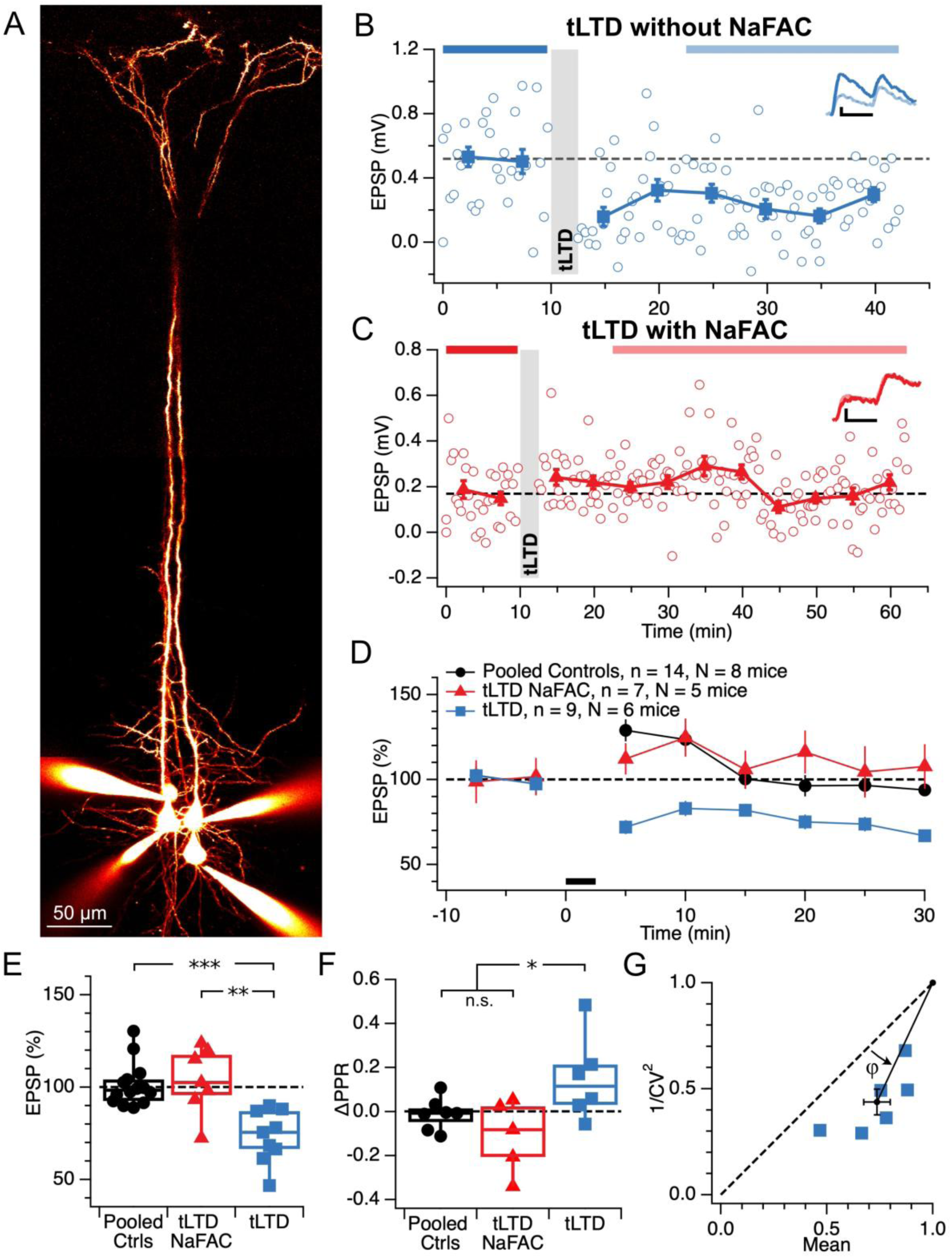
Astrocyte function is required for tLTD at L5 PC → PC synapses. **(A)** Representative quadruple whole-cell recording used to probe monosynaptic L5 PC → PC connections in acute slices. Neurons were filled with Alexa 594 for visualization. **(B)** Example tLTD experiment in a control slice. Gray bar indicates the tLTD induction period. Inset: representative averaged EPSPs before and after induction. Scale bars: 25 ms, 0.1 mV. **(C)** Example experiment in a NaFAC-treated slice showing abolished tLTD. Induction protocol and format as in (B). **(D)** Time course of normalized EPSP amplitudes from pooled control recordings (black), tLTD in NaFAC-treated slices (red), and tLTD in untreated slices (blue). Black bar: induction period. **(E)** Magnitude of plasticity across groups demonstrates that NaFAC abolished tLTD (after/before Pooled Ctrls 101 ± 3%, tLTD NaFAC 104 ± 7%, tLTD 74 ± 5%; ANOVA p < 0.001). Boxplots show medians and quartiles; whiskers denote extremes. **(F)** Changes in paired-pulse ratio (ΔPPR) were consistent with presynaptically expressed tLTD (Sjöström et al., 2003). ANOVA p < 0.05. ΔPPR of controls (−0.01 ± 0.03) and tLTD NaFAC (−0.11 ± 0.07) were indistinguishable (t-test p = 0.23) and were therefore pooled for comparison with tLTD (0.15 ± 0.08, t-test p < 0.05). **(G)** Coefficient of variation (CV) analysis supported a presynaptic locus of expression for tLTD (φ = 21 ± 3°; Wilcoxon signed rank p < 0.05) (Brock et al., 2020).

NaFAC-treated recordings showed stable responses (after/before: 103 ± 4%, *n* = 6), comparable to control stability recordings without NaFAC (100 ± 5%, *n* = 8, *p* = 0.61; pooled in Fig. 1D, E), suggesting that acute synaptic transmission was not directly impaired. However, overall experimental success was lower in NaFAC-treated slices (42% inclusion rate of recordings) than in controls (65%), suggesting possible adverse effects on slice viability.

In untreated slices, we next examined the locus of tLTD expression using PPR and CV analyses (Brock et al., 2020). A positive ΔPPR indicates decreased release probability, consistent with a presynaptic locus of expression. Similarly, because normalized 1/CV^2^ scales with release probability, a reduction in 1/CV^2^ also supports decreased presynaptic release (Brock et al., 2020). Consistent with these criteria, both PPR and CV analyses pointed to a presynaptic expression of tLTD (Fig. 1F, G), in agreement with previous reports (Sjöström et al., 2003).

These findings suggested that astrocytes may be required for L5 PC → PC tLTD, although we could not rule out potential off-target effects of NaFAC, including neuronal toxicity.

### Abolishing astrocyte Ca^2+^ signaling disrupts tLTD

To further investigate the role of astrocytes in L5 PC → PC tLTD, as suggested by the NaFAC experiments, we examined whether impairing astrocyte Ca^2+^ signaling would alter plasticity expression. We selectively expressed the human plasma membrane Ca^2+^ ATPase isoform 2w/b (hPMCA2w/b, also known as CalEx) in astrocytes by injecting AAV-pZac2.1-GfaABC1D-mCherry-hPMCA2w/b (Yu et al., 2018) into neonatal visual cortex. CalEx constitutively exports cytosolic Ca^2+^, thereby disrupting intracellular Ca^2+^ dynamics (Yu et al., 2018; Institoris et al., 2022; Guayasamin et al., 2025). Experiments were performed at P15 – P22, when astrocyte CalEx expression was robust (Fig. 2A, B).

**Figure 2.**
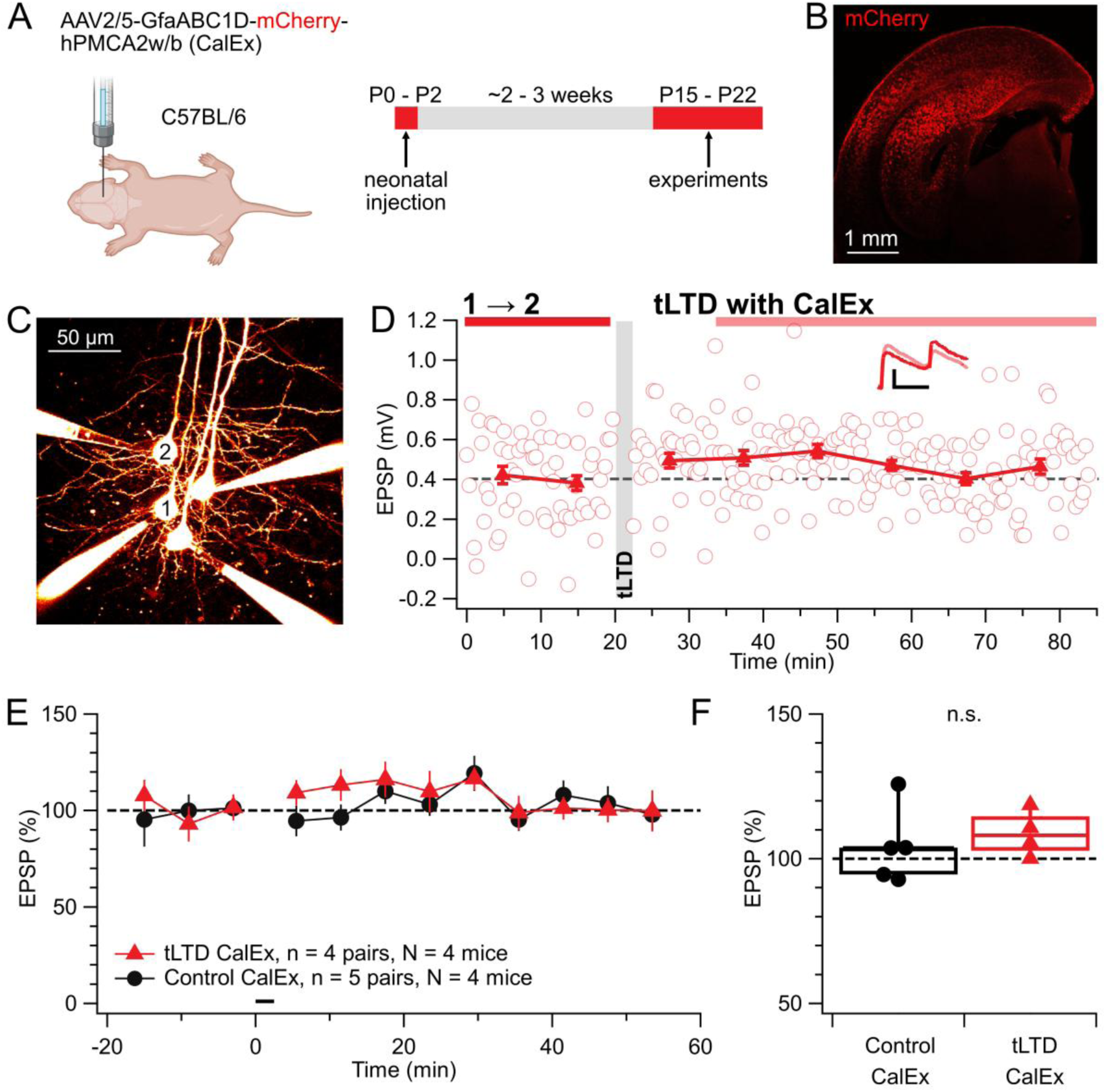
tLTD relies on astrocyte Ca^2+^ signaling. **(A)** Neonatal AAV injections to express CalEx (AAV2/5-GfaABC1D-mCherry-hPMCA2w/b) were performed at P0–P2 in C57BL/6 mice. Electrophysiology was conducted at P15 – P22. **(B)** Sample expression of CalEx in visual cortex, visualized via mCherry fluorescence following neonatal AAV injection. **(C)** Representative quadruple whole-cell recording used to probe for monosynaptic PC → PC connections in CalEx-expressing slices. Neurons were filled with Alexa 594. Presynaptic PC1 was connected to postsynaptic PC2. **(D)** Example electrophysiological trace from the same PC1 → PC2 connection as in (C). Gray shading denotes the tLTD induction period. Inset scale bars: 25 ms, 0.2 mV. **(E)** Ensemble time course of normalized EPSP amplitudes before and after induction, comparing tLTD induction (red) and control recordings (black) in CalEx-expressing slices, showing no tLTD. **(F)** Population averages showed that plasticity was indistinguishable across the two groups (after/before Control 104 ± 6% vs tLTD CalEx 109 ± 4%; t-test p = 0.54).

Using quadruple whole-cell recordings, we identified monosynaptically connected L5 PC → PC pairs in CalEx-expressing slices (Fig. 2C). Under these conditions, the post-before-pre induction protocol failed to elicit tLTD, and EPSP amplitudes remained stable, indistinguishable from non-induction control recordings (Fig. 2D, E). These results show that astrocyte Ca^2+^ signaling is required for L5 PC → PC tLTD.

To complement the global Ca^2+^ extrusion approach, we also assessed tLTD expression while loading individual astrocytes with the Ca^2+^ chelator BAPTA, taking advantage of the fact that 20 mM BAPTA can diffuse across gap-junctions throughout the astrocyte syncytium (Watanabe et al., 2023). To facilitate the search for synaptic connections while one pipette was used to patch an astrocyte, we performed experiments using 2p optomapping (Chou et al., 2024) instead of quadruple whole-cell recording. We previously demonstrated that 2p optogenetics can be reliably used to study long-term plasticity (Chou et al., 2025).

Neonatal Emx1^Cre/Cre^ mice were injected with AAV9-CAG-DIO-ChroME-ST-P2A-H2B-mRuby3 in visual cortex to express ChroME opsin in excitatory cells (Fig. 3A). Slices were preincubated with sulforhodamine 101 to label astrocytes for targeted patching under 2p imaging. In L5, PCs were patched, and EPSPs were evoked by activating ChroME-expressing candidate presynaptic cells using spiral scans at 1040 nm (Fig. 3B). A neighboring astrocyte was patched 40 – 100 µm from the PC and loaded with BAPTA via the patch pipette. When tLTD was induced with astrocytic BAPTA loading, tLTD failed (Fig. 3). Together with the CalEx data, these results support the conclusion that astrocyte Ca^2+^ signaling is essential for tLTD.

**Figure 3.**
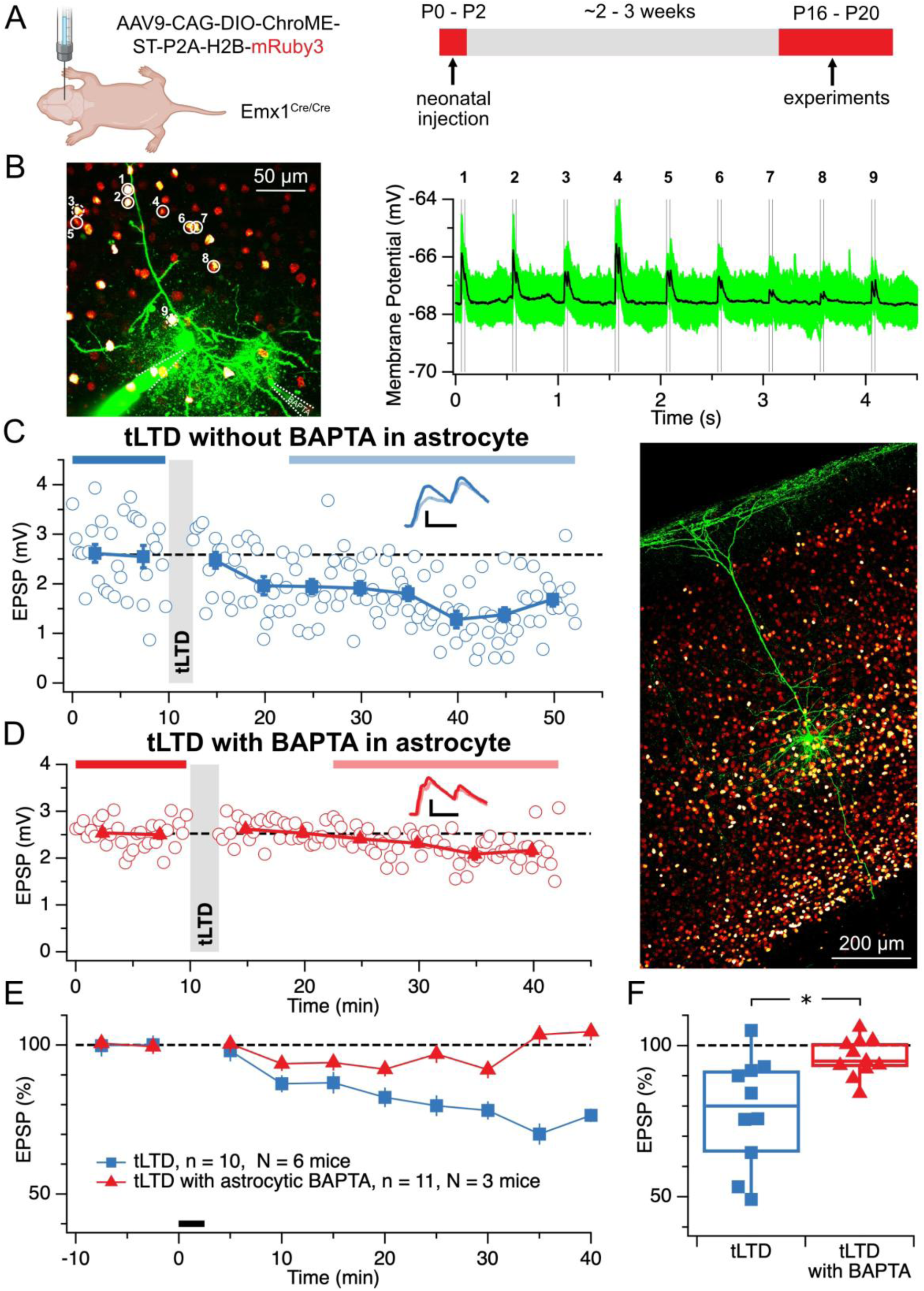
Astrocyte Ca^2+^ buffering abolishes tLTD. **(A)** ChroME was expressed in cortical excitatory neurons by neonatal viral injection (P0–P2) in Emx1^Cre/Cre^ mice. Experiments were conducted at P16–P20. **(B)** Left: experimental setup showing a patched L5 pyramidal cell (Alexa 488, green), ChroME-expressing neurons (mRuby3, red), and a nearby astrocyte patched and loaded with BAPTA (Alexa 488, green). Numbers indicate presynaptic neurons scanned via 2p spiral stimulation. Solid circles met inclusion criteria; dashed circles were excluded during analysis. Right: sample postsynaptic response traces from nine presynaptic cells. **(C)** Sample recording without astrocytic BAPTA illustrating typical amount of tLTD. Grey shading indicates induction period. Inset scale bars: 25 ms, 1 mV. **(D)** Left: sample tLTD recording with astrocytic BAPTA loading showing reduced plasticity. Induction protocol and format as in (C). Right: associated confocal image showing the biocytin-filled PC and astrocyte, with mRuby-labeled ChroME-expressing cells in red. **(E)** Ensemble averages of tLTD with or without astrocytic BAPTA. Recordings with EPSP amplitudes <0.3 mV were excluded due to poor signal-to-noise ratio. **(F)** Astrocytic BAPTA prevented tLTD (after/before tLTD 78 ± 6% vs tLTD with BAPTA 96 ± 2%; t-test p < 0.05).

### Manipulating astrocyte G-protein abolishes tLTD

Gq proteins coupled to astrocytic CB1 receptors are implicated in Ca^2+^ elevation in response to eCBs (Navarrete and Araque, 2008). We therefore asked if perturbing astrocytic Gq signaling would alter tLTD. We therefore expressed the opto-α1AR opsin (Adamsky et al., 2018) — a light-sensitive Gq-coupled receptor — in visual cortex astrocytes to enable temporally precise optogenetic manipulation (Fig. 4A).

**Figure 4.**
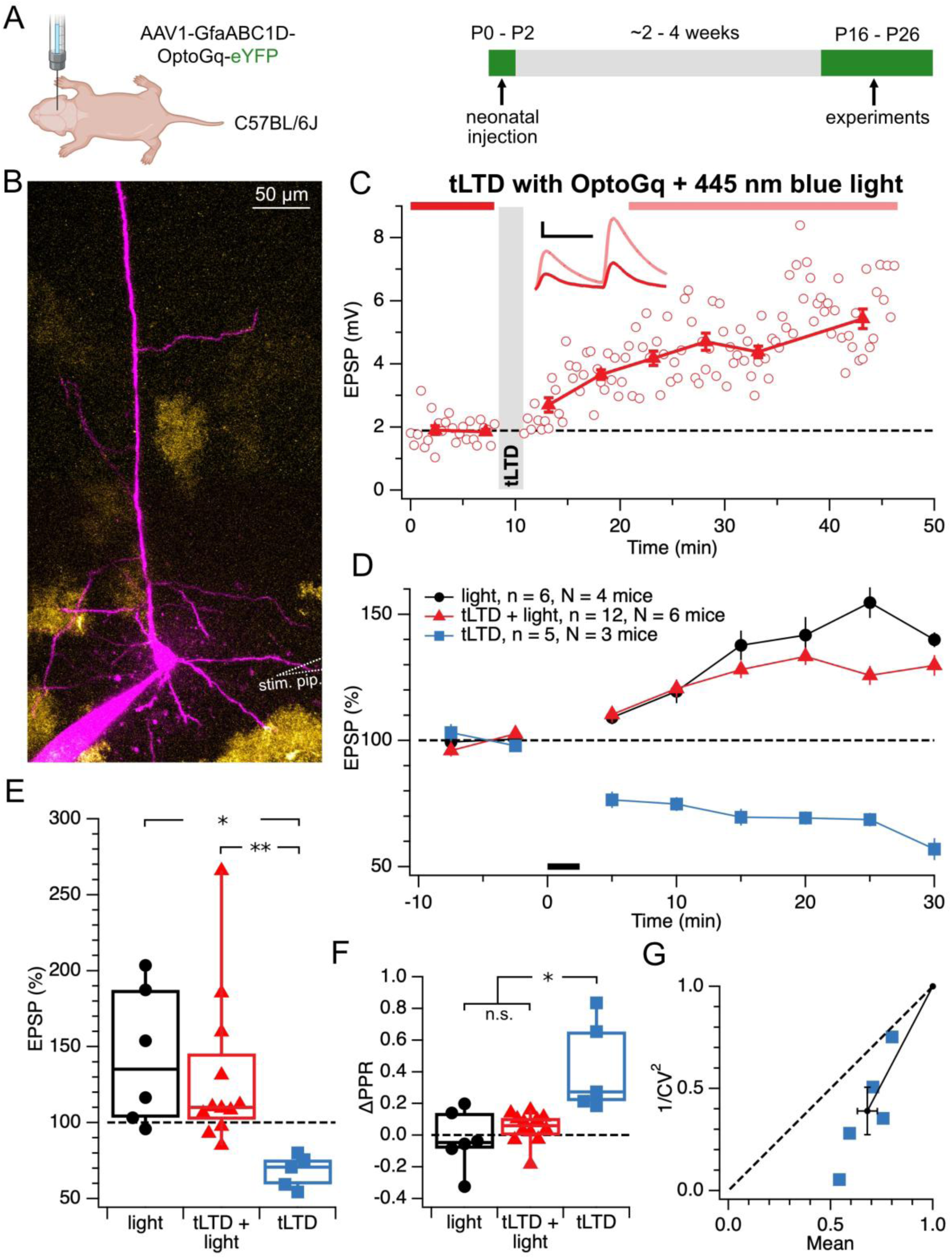
Disrupting astrocyte Gq signaling prevents tLTD. **(A)** Timeline of neonatal viral injections (P0–P2) with AAV1-GfaABC1D-OptoGq-eYFP to express a light-sensitive Gq-coupled receptor (OptoGq) in visual cortex astrocytes. Electrophysiological experiments were performed at P16–P26. **(B)** Example image showing a patched L5 pyramidal neuron filled with Alexa 594 (magenta) surrounded by eYFP-labeled astrocytes expressing OptoGq (yellow). Dotted line indicates position of extracellular stimulation pipette. **(C)** Sample recording from a tLTD + light experiment. The shaded gray bar denotes the period of plasticity induction as well as of 445-nm light delivery. Inset: representative EPSPs before and after induction. Scale bars: 25 ms, 2 mV. **(D)** Time course of normalized EPSP amplitudes for control light-only recordings (black), tLTD + light (red), and tLTD (blue). Black bar indicates induction period. **(E)** Light activation of astrocytic Gq signaling prevented tLTD and shifted plasticity toward LTP (after/before light 143 ± 18%, tLTD + light 130 ± 15%, tLTD 68 ± 5%; ANOVA p < 0.05). **(F)** ΔPPR values were consistent with presynaptically expressed tLTD (Sjöström et al., 2003). ANOVA p < 0.001. ΔPPR values in light (−0.03 ± 0.08) and tLTD + light conditions (0.05 ± 0.03) were indistinguishable (t-test p = 0.38) and were therefore pooled for comparison with tLTD (0.43 ± 0.1; t-test p < 0.05). **(G)** CV analysis in the tLTD group was consistent with a presynaptic locus of expression (φ = 16 ± 3°, Wilcoxon signed rank p = 0.06). In addition, 1⁄𝐶𝑉^2^_*norm*_ was reduced (39 ± 12%, t-test p < 0.01, n = 5), suggesting reduced release probability (Brock et al., 2020).

In acute slices from OptoGq-expressing mice, we patched L5 PCs and evoked synaptic responses using extracellular stimulation (Fig. 4B). This approach was used instead of quadruple patch or optomapping because one acquisition board was dedicated to light stimulation and spectral overlap with neuron-targeted opsins had to be avoided. During tLTD induction, 445-nm laser pulses were used to activate OptoGq in astrocytes (Fig. 4C), which altered the direction of plasticity (Fig. 4D, E). PPR and CV analyses suggested a presynaptic locus of tLTD expression (Fig. 4F, G). These results indicate that disrupting astrocytic Gq signaling abolishes tLTD.

To characterize the plasticity shift, we applied a linear mixed-effects model including initial EPSP amplitude, animal age, and sex as covariates. EPSP changes in the tLTD + light and light-only conditions were statistically indistinguishable (p = 0.93, Supp. Fig. 1). The pooled light group showed a negative correlation between EPSP change and initial amplitude (Supp. Fig. 1A). CV analysis (Supp. Fig. 1B) indicated a mixed pre- and postsynaptic locus of expression. The negative correlation between EPSP change and initial amplitude (Debanne et al., 1999; Sjöström et al., 2001; Hardingham et al., 2007), along with the CV analysis suggesting a mixed pre- and postsynaptic locus of expression (Sjöström et al., 2007), is characteristic of LTP. Taken together, these data suggest that activation of astrocytic Gq signaling prevents tLTD and instead promotes LTP.

### ACEA induces LTD at L5 PC → PC synapses via astrocyte Ca^2+^ signaling

In our previously proposed working model of tLTD (Sjöström et al., 2003), plasticity induction requires activation of presynaptic CB1 receptors by retrograde eCB signaling in conjunction with presynaptic spiking. Consistently, LTD at L5 PC → PC synapses could be induced by bath application of the synthetic CB1 receptor agonist ACEA, which mimicked and occluded tLTD (Sjöström et al., 2003).

We replicated this ACEA-induced LTD in control slices (Fig. 5A, B). In contrast, when ACEA was applied in slices expressing CalEx selectively in astrocytes, LTD was abolished (Fig. 5C-E). The loss of ACEA-induced LTD was observed only in mice older than P20, likely reflecting the time required for sufficient CalEx expression.

**Figure 5.**
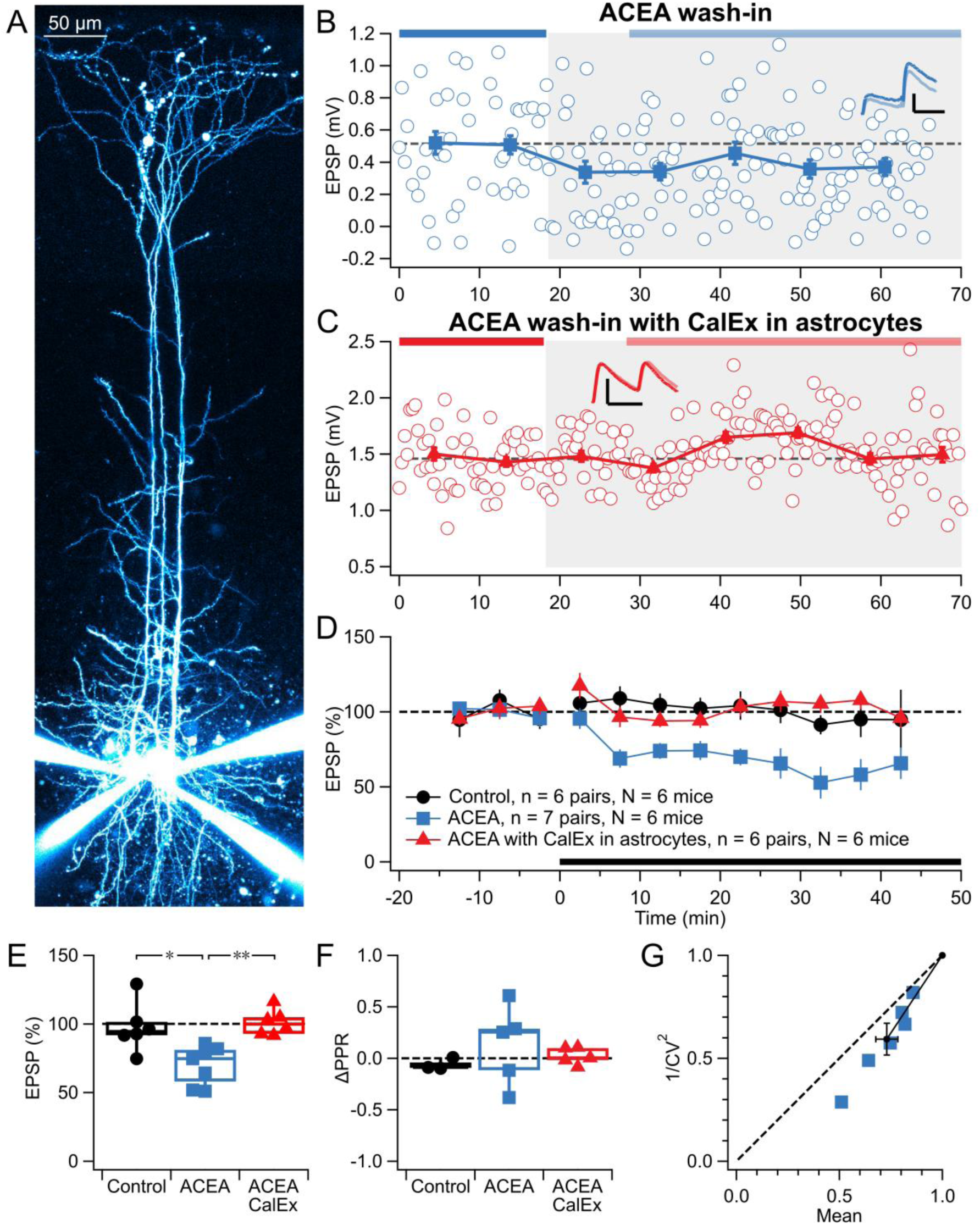
Astrocyte Ca^2+^ signaling is needed for eCB-mediated L5 PC → PC LTD. **(A)** Representative quadruple whole-cell patch recording used to probe monosynaptically connected L5 PC → PC connections in slices from control and CalEx-injected mice. Neurons were filled with Alexa 594. **(B)** Sample experiment showing a reduction in EPSP amplitude following ACEA wash-in (grey) in a control slice. Inset scale bars: 0.5 mV, 25 ms. **(C)** Sample experiment showing no EPSP reduction after ACEA wash-in (grey) in a slice with CalEx-expressing astrocytes. Inset scale bars: 1 mV, 25 ms. **(D)** Ensemble time course of normalized EPSP amplitudes, comparing control with no ACEA wash-in (black), ACEA wash-in in control slices (blue), and ACEA wash-in in CalEx expressing slices (red). Solid black line indicates the period of ACEA application. **(E)** Population data showed reduced EPSPs following ACEA wash-in in control slices (after/before Control 98 ± 7%, ACEA 70 ± 5%) but not in slices with astrocyte CalEx expression (101 ± 4%, ANOVA p < 0.01). **(F)** PPR did not detectably differ across groups (ΔPPR Control -0.06 ± 0.03, n = 3, ACEA 0.13 ± 0.2, n = 5, ACEA CalEx 0.02 ± 0.04, n = 5, ANOVA p = 0.6). **(G)** CV analysis of the ACEA group was consistent with presynaptic expression (φ = 11 ± 1°, Wilcoxon signed rank p < 0.05). In addition, 1⁄𝐶𝑉^2^_*norm*_ was reduced (59 ± 8%, t-test p < 0.01, n = 6), indicating reduced release probability (Brock et al., 2020).

PPR analysis revealed no detectable differences across conditions (Fig. 5F), suggesting that paired-pulse dynamics were not altered by ACEA application or CalEx expression. In contrast, CV analysis suggested that ACEA reduced release probability (Fig. 5G). Higher-frequency stimulation may more effectively deplete the readily releasable pool to reveal larger PPR changes due to ACEA.

Together, these results suggest that astrocyte Ca^2+^ signaling is necessary for cannabinoid-mediated LTD, paralleling its requirement for tLTD. These findings further support the involvement of astrocytes in L5 PC → PC tLTD.

### Astrocyte-Specific Deletion of CB1 Receptors Abolishes tLTD

Astrocytes possess CB1 receptors (Navarrete and Araque, 2010), so we tested if they are needed for L5 PC → PC tLTD. To specifically delete CB1 receptors in astrocytes, we either crossed CB1^f^ and Aldh1l1-Cre mice, or we injected visual cortex of CB1^f^ mice with AAV5-GfaABC1D-Cre-4×6T (Fig. 6A). To tag astrocytes, we co-injected a Cre-dependent EGFP reporter (Fig. 6A, B).

**Figure 6.**
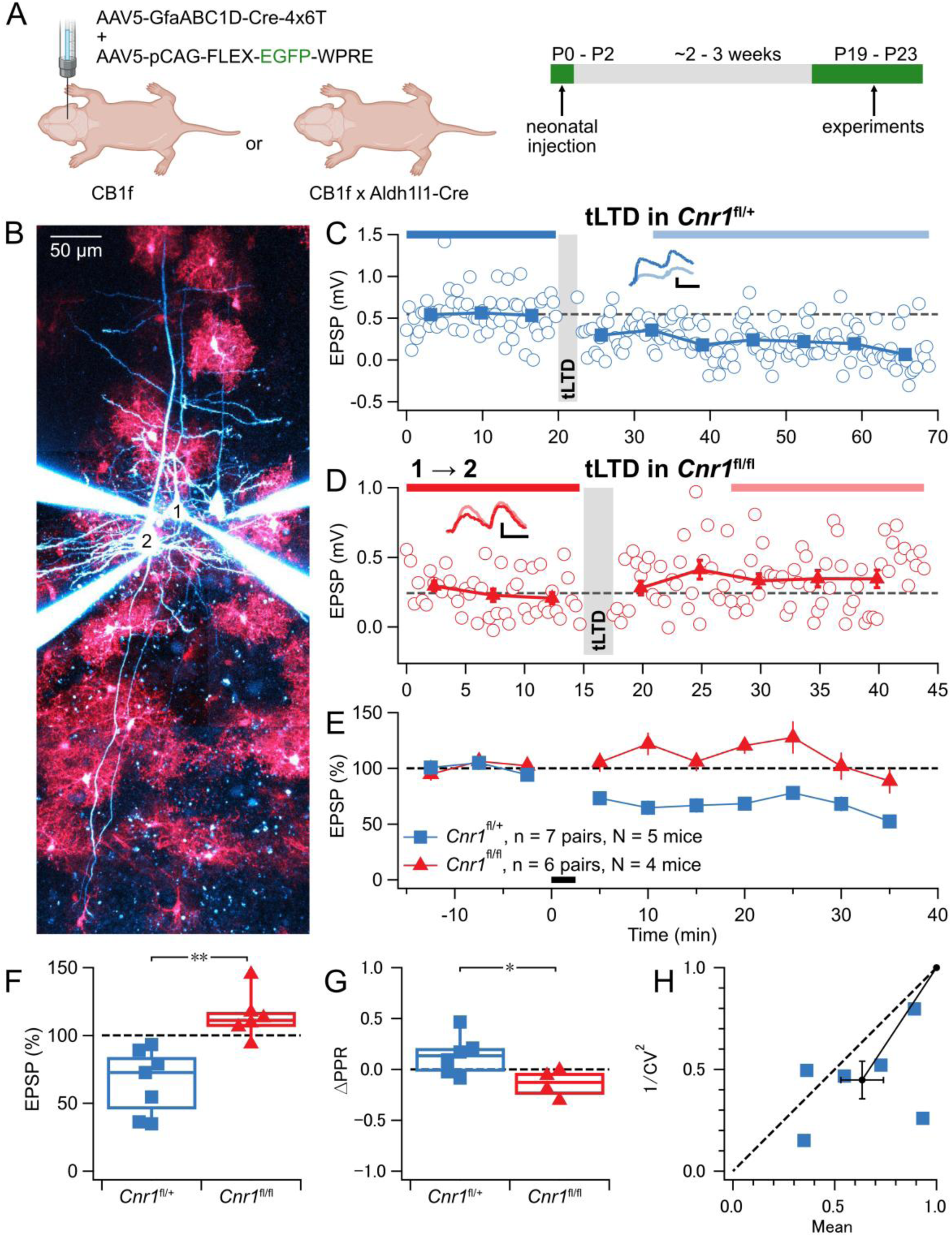
Astrocyte-specific deletion of CB1 receptors prevents L5 PC → PC tLTD. **(A)** Neonatal AAV injections were performed at P0–P2 in heterozygous or homozygous CB1^f^ mice, or CB1^f^ mice were bred with Aldh1l1-Cre mice. Experiments were conducted at P19–P23. **(B)** Representative quadruple-patch experiment used to assess monosynaptic PC → PC connections (blue, Alexa 594) in slices with astrocyte-specific *Cnr1* deletion (red, EGFP). Presynaptic PC1 was connected to postsynaptic PC2. **(C)** Example recording from tLTD induction experiments in *Cnr1*^fl/+^ slice. Gray bar indicates the tLTD induction period. Inset scale bars: 25 ms and 0.2 mV. **(D)** Example recording from the same PC1 → PC2 connection as in (B), showing failed tLTD in a *Cnr1*^fl/fl^ slice. Gray bar indicates the tLTD induction period. Inset scale bars: 25 ms and 0.2 mV. **(E)** tLTD was robust in *Cnr1*^fl/+^ (blue) but not in *Cnr1*^fl/fl^ slices (red). **(F)** Homozygous astrocyte-specific CB1 receptor deletion abolished tLTD (Cnr1^fl/fl^: 114 ± 7% vs 100%, one-sample t-test p = 0.1), whereas heterozygous deletion did not (Cnr1^fl/+^: 68 ± 9% vs 100%, one-sample t-test p < 0.05; genotype comparison t-test p < 0.01). **(G)** ΔPPR for *Cnr1*^fl/+^ group was consistent with presynaptically expressed tLTD (ΔPPR *Cnr1*^fl/+^ 0.14 ± 0.08, n = 6 vs *Cnr1*^fl/fl^ -0.14 ± 0.07, n = 4, t-test p < 0.05). **(H)** CV analysis in the *Cnr1*^fl/+^ group showed most data points below the identity line, consistent with a presynaptic expression (φ = 13 ± 6°, Wilcoxon signed rank p = 0.09). In addition, 1⁄𝐶𝑉^2^_*norm*_ was reduced (45 ± 9%, t-test p < 0.01, n = 6), suggesting reduced release probability (Brock et al., 2020).

In homozygous CB1^fl/fl^ mice for which the *Cnr1* gene was completely deleted in astrocytes, tLTD could not be induced (Fig. 6D, E, F). In contrast, heterozygous CB1^fl/+^ mice, which retained one functional copy of the *Cnr1* gene, exhibited normal tLTD (Fig. 6C, E, F). The degree of tLTD were indistinguishable when CB1 receptors in CB1^fl/+^ mice were deleted via AAV (after/before 68 ± 17%, n = 3) or when crossed with Aldh1l1-Cre mice (after/before 69 ± 12%, n = 4, t-test p = 0.96; pooled in Fig. 6E – H). PPR and CV analysis also confirmed a presynaptic locus of tLTD expression in CB1^fl/+^ mice (G, H).

These findings show that tLTD relies on astrocytic CB1 receptors, and that partial CB1 receptor deletion does not disrupt this form of plasticity. Consistent with impaired tLTD and a reduced capacity for synaptic weakening, age-matched EPSP amplitudes were furthermore larger in CB1 knockout and CalEx conditions (Supp. Fig. 2).

### CB1 receptor activation alters astrocyte Ca^2+^ dynamics

We argued that if tLTD relies on astrocytic CB1 receptor signaling, astrocytic Ca^2+^ activity may be altered by ACEA and by tLTD.

Using 2p Ca^2+^ imaging (Fig. 7A – C), we found that ACEA increased astrocyte Ca^2+^ event frequency and reduced event duration compared to control (Fig. 7D). In addition, Ca^2+^ activity became more spatially coordinated within individual astrocytes, as reflected by increased correlations across regions of interest (Fig. 7E). These changes are consistent with coordinated activation following CB1 receptor stimulation.

**Figure 7.**
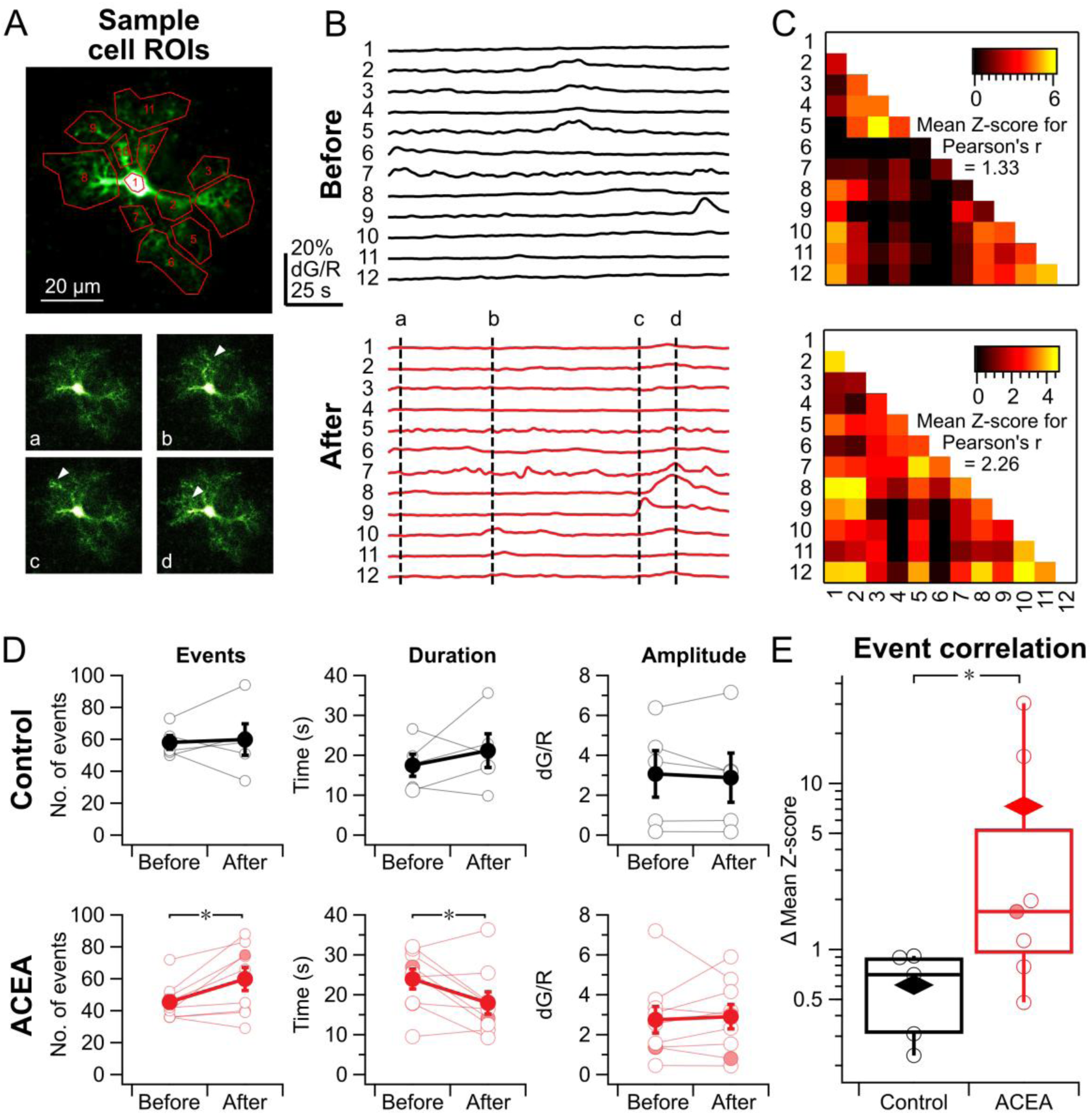
CB1 receptor activation alters astrocyte Ca^2+^ dynamics. **(A)** Fluo-5F Ca^2+^ signals (green, arrowheads) were recorded as a 150-second-long movie (see Videos S1 and S2) of this sample astrocyte. During offline analysis, twelve ROIs (red, top) were manually selected. Comparing to relatively quiet sample frame *a* (bottom), Ca^2+^ transients are visible in in sample frames *b*, *c*, and *d* (arrowheads). **(B)** Based on ROIs in (A), fluorescence was quantified as dG/R sweeps before and after ACEA wash-in. Vertical dashed lines denote the timepoints of frames *a* through *d* depicted in (A), bottom. Note synchrony across most ROIs at timepoint *d*. **(C)** Sample ROI cross-correlation matrices before and after ACEA wash-in for astrocyte in (A). For each matrix, a single mean Z-score for Pearson’s r was calculated as a metric for how correlated the activity of a cell was. **(D)** The number of Ca^2+^ events was increased by ACEA (45 ± 4 vs 60 ± 7, n = 9; paired t-test p < 0.05) but not in controls (58 ± 4 vs 60 ± 10, n = 5; p = 0.8). The duration of Ca^2+^ events, however, decreased during ACEA wash-in (24 ± 3 s vs 18 ± 3 s, n = 9; p < 0.05) but not in control (18 ± 3 s vs 21 ± 4 s, n = 5; p = 0.4). Amplitude did not differ for either condition (ACEA 2.7 ± 0.7 vs 2.9 ± 0.6, n = 9, p = 0.7, and Control 3.1 ± 1 vs 2.9 ± 1, n = 5, p = 0.6). Closed circles indicate sample astrocyte shown in (A). **(E)** ACEA increased event correlation (Δ mean Z-score: 7.3 ± 4, n = 7) compared to controls (0.6 ± 0.1, n = 5; Mann-Whitney U test p < 0.05). Diamonds denote means. Closed circle indicates sample astrocyte shown in (A).

However, in astrocytes patched <100 µm from neurons that underwent tLTD induction, there were no detectable changes in Ca^2+^ event frequency (before: 63 ± 14 vs after: 70 ± 9 , n = 3; paired t-test p = 0.5) or duration (21 ± 9 s vs 16 ± 6 s , n = 3; paired t-test p = 0.5). In addition, Ca^2+^ activity did not synchronize, as reflected by unchanged cross correlation across regions of interest (Δ mean Z-score for tLTD: 0.6 ± 0.2, n = 3 vs Control: 1.4 ± 0.5, n = 3; t-test p = 0.1).

This contrasts with previous reports that astrocyte Ca^2+^ activity increases during tLTD induction (Min and Nevian, 2012) and indicates that, under our conditions, tLTD does not measurably alter astrocyte Ca^2+^ dynamics. However, this does not exclude the possibility that synapse-specific astrocyte Ca^2+^ microdomains were engaged during tLTD induction but were too small, too brief, or too spatially restricted to be resolved under our recording conditions.

In conclusion, CB1 receptor activation robustly alters astrocyte Ca^2+^ dynamics, whereas tLTD induction does not produce detectable changes under our experimental conditions.

## Discussion

Here, we revisited the cellular basis of tLTD at visual cortex L5 PC → PC synapses. Although this plasticity has been attributed to presynaptic NMDA and CB1 receptors, our findings reveal that astrocytes are a critical third player. Using complementary loss-of-function approaches, we found that astrocyte Ca^2+^ signaling and astrocytic CB1 receptors are essential for tLTD induction. Moreover, optogenetic activation of astrocytic Gq signaling during induction abolished tLTD and shifted the outcome toward potentiation, highlighting a bidirectional astrocytic influence.

### Astrocytes are required for L5 PC → PC tLTD

We found that astrocytes are essential for inducing L5 PC → PC tLTD in mouse visual cortex. Disruption of glial metabolism with NaFAC — a glia-specific Krebs cycle inhibitor (Swanson and Graham, 1994) — abolished tLTD, consistent with prior reports that NaFAC impairs both LTD (Zhang et al., 2008) and LTP (Fossat et al., 2012; Han et al., 2015; Boddum et al., 2016). To rule out nonspecific effects, we also employed more targeted astrocyte-specific approaches.

We interfered with astrocyte Ca^2+^ signaling using two orthogonal strategies: the plasma membrane Ca^2+^ ATPase CalEx (Yu et al., 2018), and intracellular loading of the fast Ca^2+^ chelator BAPTA. Both manipulations reliably abolished tLTD, confirming that astrocyte Ca^2+^ transients are required. BAPTA is expected to target fast, local astrocyte Ca^2+^ signals (Tsien, 1980), while CalEx dampens global Ca^2+^ transients in astrocytes by extruding cytosolic Ca^2+^ (Yu et al., 2018). Their convergent effects suggest that both local and global astrocytic Ca^2+^ signals contribute to tLTD.

We found that buffering Ca^2+^ in astrocytes within ∼100 µm of the postsynaptic neuron was sufficient to abolish tLTD, consistent with BAPTA spreading via gap junctions throughout the syncytial astrocyte network (Watanabe et al., 2023). This outcome agrees with prior studies in cortex and hippocampus (Andrade-Talavera et al., 2016; Andrade-Talavera et al., 2024; Martínez-Gallego et al., 2024).

Our findings thus revise our earlier model of L5 PC → PC tLTD, which posited that coincident activation of presynaptically located CB1 and NMDA receptors drove tLTD (Sjöström et al., 2003; Duguid and Sjöström, 2006). Instead, our new results position astrocytes as critical intermediaries in this form of STDP (Fig. 8).

**Figure 8.**
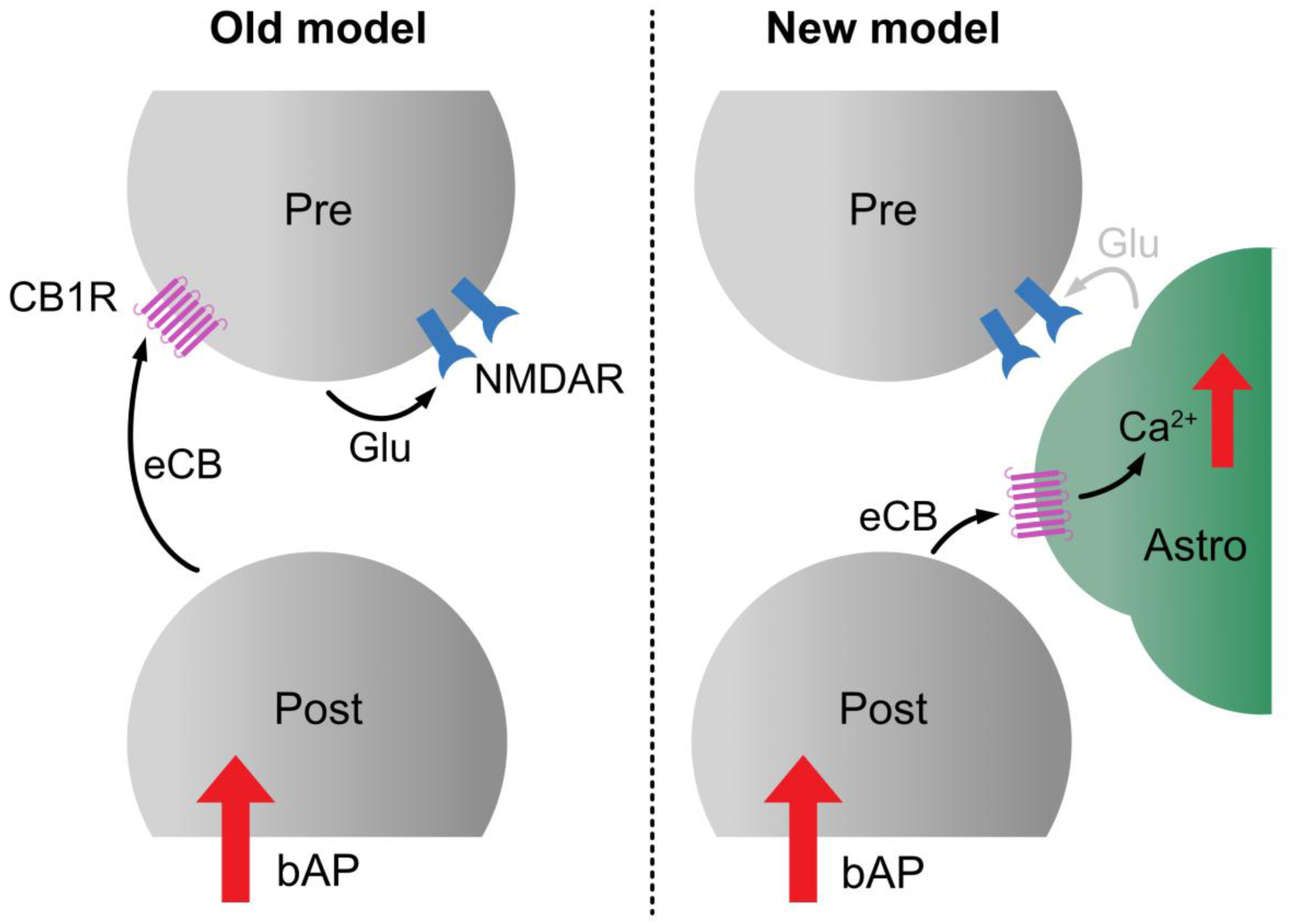
Updated working model of L5 PC → PC tLTD. Left: The original model of tLTD proposed that coincident activation of presynaptic NMDA and CB1 receptors is required to induce tLTD at L5 PC → PC synapses (Sjöström et al., 2003). Right: We now update this framework to include a central role for astrocytes, in which astrocytic CB1 receptors and Ca^2+^ signaling mediate the induction of tLTD. While in L4 → L2/3 tLTD requires glutamate release from astrocytes (Min and Nevian, 2012), tLTD at other synapses requires different gliotransmitters, such as D-serine (Andrade-Talavera et al., 2016; Andrade-Talavera et al., 2024; Martínez-Gallego et al., 2024). Pre: presynaptic terminal, Post: postsynaptic terminal, Astro: astrocyte, bAP: back propagating action potential, eCB: endocannabinoid, CB1R: cannabinoid receptor type-1, NMDAR: NMDA receptor, Glu: glutamate.

### Astrocytic Gq signaling promotes potentiation

Optogenetic activation of astrocytic Gq signaling disrupted tLTD and drove what appeared to be LTP, as evidenced by increased EPSP amplitude, an inverse correlation with baseline strength (Sjöström et al., 2001), and a shift in CV consistent with mixed pre- and postsynaptic expression (Sjöström et al., 2007). These observations suggest that astrocytes not only gate plasticity but can shift its polarity.

It is possible that Gq signaling biases gliotransmitter release toward LTP-promoting cues such as D-serine or glutamate. Astrocyte-driven LTP following Gq-DREADD activation has been reported to be NMDA receptor-dependent (Adamsky et al., 2018). Our results are broadly consistent with this and additionally suggest that astrocytes, via Gq activation, can override tLTD and instead promote potentiation.

This effect appears to be context- and circuit-dependent. While astrocyte Gq activation triggered LTP at our visual cortex L5 PC → PC synapses, Gq activation induces LTD at corticostriatal synapses (Cavaccini et al., 2020), highlighting the specificity of astrocyte-mediated modulation. Other modes of astrocyte activation — including melanopsin stimulation (Mederos et al., 2019) and cAMP elevation (Zhou et al., 2021) — can also induce LTP. Multiple astrocytic signaling pathways may thus converge to facilitate potentiation.

Together, these findings expand current models of STDP by showing that astrocytes are not passive but active integrators in plasticity.

### Astrocyte CB1 receptors mediate cannabinoid-dependent tLTD

We identified astrocyte CB1 receptors as an essential locus of eCB signaling in tLTD. Bath application of ACEA, a synthetic analog of anandamide, mimicked and occluded tLTD (Sjöström et al., 2003). This aligns with recent findings that anandamide preferentially activates astrocytic CB1Rs, while 2-arachidonoylglycerol preferentially engages neuronal CB1Rs (Noriega-Prieto et al., 2025). The sensitivity of L5 PC → PC synapses to ACEA, along with the abolition of tLTD by astrocyte-targeted CalEx, suggests that the relevant eCB signaling is mediated by astrocytes rather than neurons.

Crucially, conditional deletion of CB1 receptors from astrocytes abolished tLTD, similar to disrupting astrocytic Ca^2+^ signaling. Astrocytes thus decode activity-dependent eCB release via their CB1 receptors that could be Gq-coupled (Navarrete and Araque, 2008), mobilize intracellular Ca^2+^, and communicate back to neurons through the release of gliotransmitters, functioning as critical intermediaries between pre- and postsynaptic compartments.

Both deletion of CB1 receptors from astrocytes and disruption of astrocytic Ca^2+^ signaling resulted in increased strength of L5 PC → PC synapses. This is consistent with impaired tLTD and a reduced capacity for synaptic weakening, although contributions from other disruptions of astrocyte function cannot be excluded.

Astrocytic CB1 receptors are increasingly recognized as modulators of plasticity and behavior. They have been implicated in D-serine regulation and recognition memory (Robin et al., 2018), in stress resilience (Dudek et al., 2025), and in context-dependent inflammatory signaling (Colomer et al., 2026). The latter study proposed that distinct astrocytic CB1 receptor pools located at the plasma membrane or within mitochondria may mediate opposing effects depending on the inflammatory milieu.

Our findings add to this growing body of work, highlighting that CB1 receptor-centric plasticity models not involving astrocytes may need to be revisited.

### Functional implications of tripartite tLTD

Our findings place astrocytes as key contributors to cortical tLTD, suggesting a functional tripartite L5 PC → PC synapse (Perea et al., 2009). This supports three-factor learning rules (McFarlan et al., 2023), where a modulatory signal influences Hebbian plasticity (Pawlak et al., 2010). Astrocytes are ideally suited for this third role, integrating local activity and neuromodulatory context to gate plasticity outcomes (Falcón-Moya et al., 2020; Martinez-Gallego et al., 2022; Lefton et al., 2025).

Astrocyte involvement may help explain why STDP varies across synapse types (Larsen and Sjöström, 2015; McFarlan et al., 2023). Differences in astrocyte coupling, receptor profiles, and brain state sensitivity provide a flexible substrate for context-dependent circuit tuning (Min and Nevian, 2012; Andrade-Talavera et al., 2016; Letellier et al., 2016; Falcón-Moya et al., 2020; Letellier and Goda, 2023).

Moreover, astrocyte calcium dynamics track global states like arousal and sleep (Poskanzer and Yuste, 2016). Tripartite tLTD thus enables participation in behaviorally relevant circuit reconfiguration. By identifying astrocytes as indispensable for tLTD, our work reframes this plasticity as a tripartite learning process.

### Caveats and limitations

This study was conducted in acute slices from juvenile mice. It remains to be determined whether our findings generalize to more mature developmental stages or in vivo conditions.

The OptoGq experiments relied on extracellular stimulation, which in neocortex may non-specifically recruit multiple synaptic pathways (Dragsted et al., 2025). Given the well-established synapse-type specificity of cortical plasticity (Larsen and Sjöström, 2015; McFarlan et al., 2023), such non-specificity could confound interpretation. However, the majority of our experiments targeted identified L5 PC → PC connections, either using paired recordings (Lalanne et al., 2016) or 2p optogenetic stimulation (Chou et al., 2024). This allowed us to link observed plasticity effects specifically to the L5 PC → PC synapse type.

Astrocyte Ca^2+^ imaging was acquired at 2.11 Hz, sufficient to capture global Ca^2+^ transients in processes, but likely insufficient to detect fast, localized microdomain events in fine astrocytic branches. Functionally important Ca^2+^ signals may therefore have gone undetected.

While PPR and CV analyses suggested a presynaptic locus of expression, these are indirect measures (Brock et al., 2020). The downstream pathways through which astrocytic CB1 receptor activation modulates presynaptic function remain unclear and warrant further investigation.

### Conclusions and outlook

Our study challenges canonical neuron-centric models of synaptic plasticity by demonstrating that astrocytes decode eCB signaling to enable tLTD (Fig. 8). By identifying astrocytes as active participants, our results support the emerging tripartite framework in which glia dynamically gate synaptic change (Perea et al., 2009; Henneberger et al., 2010). More broadly, our findings place astrocytes within three-factor learning rules, in which modulatory signals shape synaptic plasticity (Pawlak et al., 2010; Gerstner et al., 2018; McFarlan et al., 2023) and add to the growing body of work showing the involvement of astrocytes in STDP (Min and Nevian, 2012; Andrade-Talavera et al., 2016; Andrade-Talavera et al., 2024; Martínez-Gallego et al., 2024).

The finding that astrocytic Gq signaling controls synaptic plasticity suggests that astrocytes are local arbiters of synaptic modification, potentially linking it to network state or neuromodulatory context (Poskanzer and Yuste, 2016). Such mechanisms may add computational power, for instance to address the credit assignment problem.

Future work will clarify how astrocytic CB1 receptor activation modulates synaptic function, determine the gliotransmitters involved, and test whether astrocyte-dependent tLTD operates in vivo. Given links to eCB signaling and glial dysfunction in disease, these insights may inform future therapies targeting circuit pathology.

## Acknowledgements

We thank Inbal Goshen and her lab for the generous gift of OptoGq AAV. We thank Alanna Watt, Keith Murai, Aparna Suvrathan, Anne McKinney, Jonathan Britt, Alfonso Araque and his lab, Sabine Rannio, Elia Painchaud-Lakatos, Meghan Appleby, and Mackenzie Fredericks for help and insights. We thank Christina Chou for guidance on optomapping, Jialin Ding and Zoe Harrington for help with genotyping, staining, and imaging. We thank all other members of the Sjöström lab for support and useful discussions. We thank the Molecular Imaging Platform at the Research Institute of the McGill University Health Centre and facility staff for their contribution to this publication. Neonatal injection schematic in Fig. 2, 3, 4, 6 was created in BioRender. Watanabe, A. (2026) https://BioRender.com/ipt45k5

## Author Contribution

AW, CG, and PJS conceptualized the study and designed the experiments. AW and CG carried out experiments. AW, CG, SA, and PJS conducted formal analysis. PJS wrote custom software. AW, CG and PJS wrote the manuscript.

## Data Availability Statement

The raw data supporting the conclusions of this manuscript will be made available by the authors, without undue reservation, to any qualified researcher.

