## Supplemental data for "Layer-5 Pyramidal Cell tLTD Requires Astrocytic Ca^2+^ and CB1 Receptor Signaling"

### Supplementary Figures

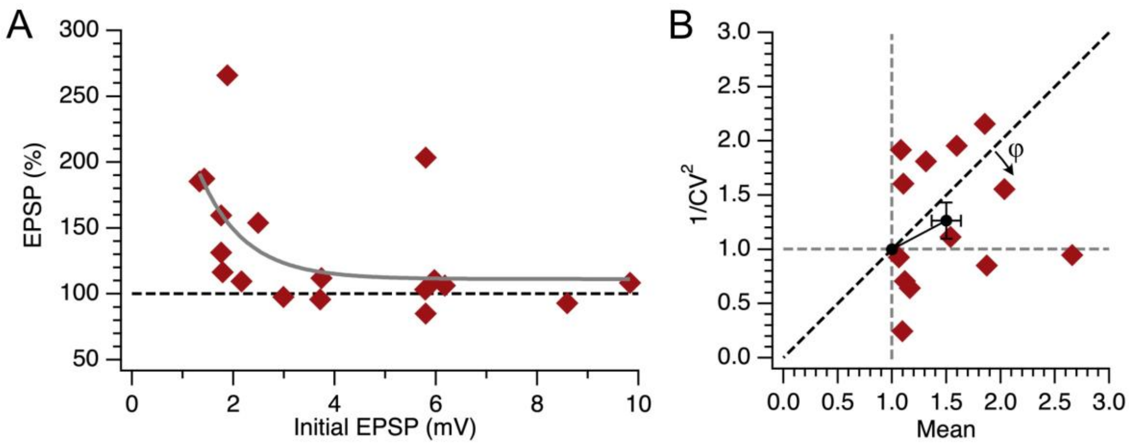


***Supplementary Figure 1 | Astrocyte Gq activation potentiated L5 PC → PC synapses with features consistent with LTP.***

**(A)** Weak responses potentiated more than strong ones ($p < 0.05$, $n = 18$), meaning LTP saturated for sufficiently strong synapses, as previously demonstrated for L5 PC → PC LTP in the rat (Sjöström et al., 2001) as well as for other synapse types (Liao et al., 1992; Bi and Poo, 1998; Debanne et al., 1999; Montgomery et al., 2001; Hardingham et al., 2007; Zhang and Oertner, 2007). For the pooled light (light and tLTD + light) group, after/before percentage EPSP was plotted against the initial EPSP amplitude, with grey line denoting the exponential fit. Statistical analysis was done using a linear mixed model incorporating age and sex.

**(B)** CV analysis indicated a mixed pre- and postsynaptic locus of LTP expression ($\varphi = 37 \pm 17°$, Wilcoxon signed rank $p = 0.08$), as previously shown for L5 PC → PC LTP in the rat (Sjöström et al., 2007), although we found no evidence that release probability systematically increased ($1/CV_{norm}^2$ = 126% ± 17%, $n = 13$, t-test $p = 0.14$). Only recordings with at least 5% plasticity were included in this analysis.

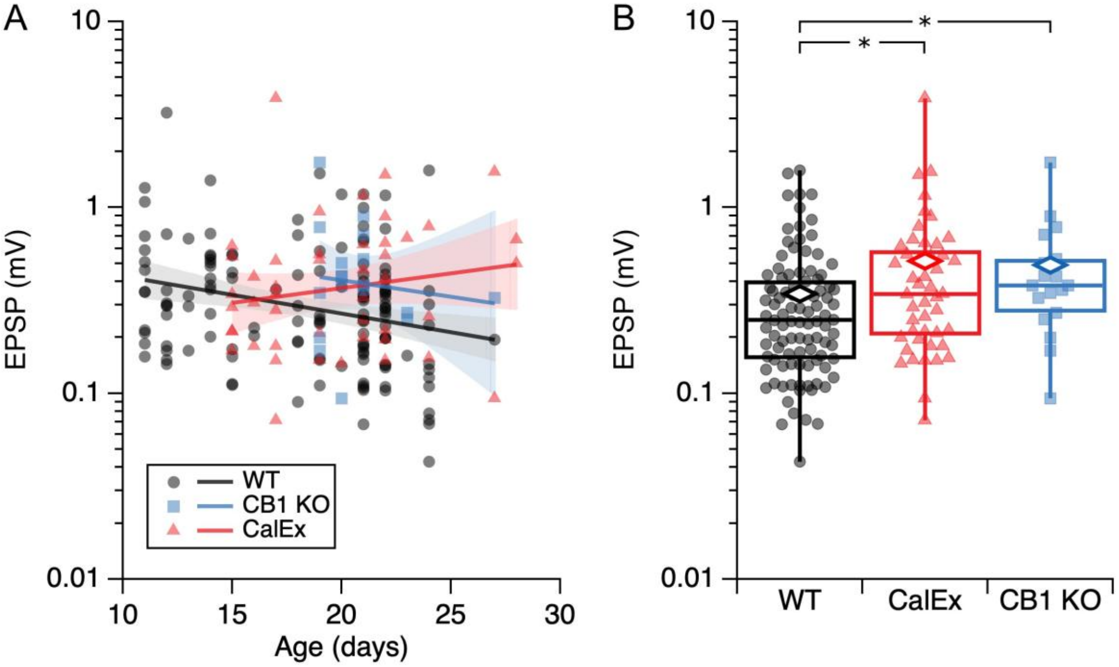


***Supplementary Figure 2 | Synaptic weight depends on age and on astrocyte signaling***

**(A)** Over age, PC → PC EPSP amplitude in WT mice decreased (log-transformed data; $\beta = -0.046$, $p < 0.01$ over age, $n = 139$), as previously reported (Oswald and Reyes, 2008). However, in CalEx, synaptic strength showed a different trend with age ($\beta = 0.037$, $p < 0.05$ vs WT, $n = 47$), consistent with CalEx expression requiring time to take effect. For CB1 KO, synaptic weight change over age did not detectably differ from WT ($\beta = 0.041$, $p = 0.99$ vs WT). Statistical analysis was done using a linear mixed model (Methods). WT data included L5 PC → PC paired recordings pooled with L5 PC → PC optomapping data from Chou et al. (2024).

**(B)** Age-matched PC → PC EPSP amplitudes were larger in CalEx expressing and in CB1 deletion mice than in WT mice (Kruskal-Wallis $p < 0.01$; WT: mean ± SD = 0.34 ± 0.03 mV, $n = 100$; CalEx: 0.52 ± 0.09 mV, $n = 47$; CB1 KO: 0.49 ± 0.09 mV, $n = 18$; Mann-Whitney U tests pairwise), consistent with tLTD being abolished by these manipulations. Age matching was achieved by selecting EPSP amplitudes from animals aged ≥ P15, which rendered ages indistinguishable (Kruskal-Wallis $p = 0.23$; all pairwise Mann-Whitney $p > 0.05$). Boxplots show medians and quartiles, with whiskers denoting extrema. Diamonds denoting the means are characteristically larger than the medians for long-tailed EPSP amplitude distributions (Song et al., 2005; Chou et al., 2024), suggesting non-parametric statistical comparison.
